# *Staphylococcus aureus* α-toxin induces acid sphingomyelinase release from a human endothelial cell line

**DOI:** 10.1101/2021.03.29.437456

**Authors:** David Krones, Marcel Rühling, Katrin Anne Becker, Tobias C. Kunz, Carolin Sehl, Kerstin Paprotka, Erich Gulbins, Martin Fraunholz

**Affiliations:** Chair of Microbiology, Biocenter, University of Würzburg, Würzburg, Germany; Institute of Molecular Biology, University of Duisburg-Essen, University Hospital, Essen, Germany; Department of Surgery, University of Cincinnati, Collage of Medicine, Cincinnati, USA

## Abstract

*Staphylococcus aureus* (*S. aureus*) is well known to express a plethora of toxins of which the pore-forming hemolysin A (α-toxin) is the best-studied cytolysin. Pore-forming toxins (PFT) permeabilize host membranes during infection thereby causing concentration-dependent effects in host cell membranes ranging from disordered ion fluxes to cytolysis. Host cells possess defense mechanisms against PFT attack, resulting in endocytosis of the breached membrane area and delivery of repair vesicles to the insulted plasma membrane as well as a concurrent release of membrane repair enzymes. Since PFTs from several pathogens have been shown to recruit membrane repair components, we here investigated whether staphylococcal α-toxin is able to induce these mechanisms in endothelial cells. We show that *S. aureus* α-toxin induced increase in cytosolic Ca^2+^ in endothelial cells, which was accompanied by p38 MAPK phosphorylation. Toxin challenge led to increased endocytosis of an extracellular fluid phase marker as well as increased externalization of LAMP1-positive membranes suggesting that peripheral lysosomes are recruited to the insulted plasma membrane. We further observed that thereby the lysosomal protein acid sphingomyelinase (ASM) was released into the cell culture medium. Thus, our results show that staphylococcal α-toxin triggers mechanisms in endothelial cells, which have been implicated in membrane repair after damage of other cell types by different toxins.

## Introduction

The opportunistic pathogen *Staphylococcus aureus* (*S. aureus*) produces a plethora of toxins of which the pore-forming α-toxin is most investigated (e.g. reviewed in Bhakdi and Tranum-Jensen, 1991; von Hoven et al., 2019). However, the bi-component PFTs γ-hemolysin and the so called leukocidins also form pores in target membranes, but are specific for certain cell types or even species-specific (Alonzo and Torres, 2014), whereas α-toxin is able to permeabilize different cell types (Berube and Bubeck Wardenburg, 2013). Further, the so-called phenol-soluble modulins (PSM) are secreted by *S. aureus* in copious amounts (Wang et al., 2007). PSMs, membrane active peptides, are strongly cytolytic in absence of serum lipoproteins (Surewaard et al., 2012) and thus are thought to be required for the intracellular virulence of *S. aureus* upon internalization by epithelial or endothelial cells, where they can cause the translocation of the pathogen from phago-endosomes to the host cytoplasm (Grosz et al., 2014).

*S. aureus* α-toxin induces cell death in mammalian host cells (Menzies and Kourteva, 2000; Haslinger-Loffler et al., 2005). In sensitive target cell membranes, α-toxin monomers assemble to a heptameric pre-pore (Bhakdi and Tranum-Jensen, 1991) and subsequent pore formation can lead to calcium influx and efflux of potassium (Fink et al., 1989; Walev et al., 1993; Kwak et al., 2012), or even larger molecules such as ATP (Gierok et al., 2014), eventually leading to host cell death (Baaske et al., 2016). Specific binding of α-toxin to *A Disintegrin And Metalloproteinase Domain-containing protein* 10 (ADAM10) facilitates pre-pore assembly on target membranes (Popov et al., 2015). Furthermore, the activated metalloprotease leads to degradation of cell barrier proteins in epithelial and endothelial cells, allowing for *S. aureus* deep tissue penetration (Inoshima et al., 2011; Becker et al., 2018). α-toxin sensitivity of target membranes is further dependent on sphingomyelin, since mutants in sphingomyelin synthetase SGMS1 were identified as α-toxin resistant in a genome-wide CRISPR screen (Virreira Winter et al., 2016) and treatment with the bacterial sphingomyelinase (bSMase) staphylococcal β-toxin abrogated α-toxin sensitivity in epithelial cells HBE16o^-^ (Ziesemer et al., 2019).

Mammalian cells also express sphingomyelinases, which serve crucial cellular functions in lipid signaling, membrane fusion, apoptosis induction (Santana et al., 1996; Zhao et al., 2016), infection (Hauck et al., 2000; Riethmüller et al., 2006; Grassmé et al., 2008; Faulstich et al., 2015; Seitz et al., 2015; Peters et al., 2019) or membrane repair mechanisms after plasma membrane insult, e.g. by pore forming toxins such as streptolysin O (SLO) or listeriolysin O (LLO) (Andrews et al., 2014). Bacteria (Grassmé et al., 1997; Grassmé et al., 2003; Simonis and Schubert-Unkmeir, 2018), viruses (Grassmé et al., 2005; Miller et al., 2012; Shivanna et al., 2015) as well as fungi (Bryan et al., 2015) have been shown to exploit enzymes of the sphingolipid metabolism. Upon *S. aureus* infections, a rapid activation of acid sphingomyelinase (ASM) occurs and this activation is triggered by staphylococcal α-hemolysin (Ma et al., 2017; Becker et al., 2018).

Plasma membrane damage by PFT may trigger membrane repair mechanisms, which depend on the pore size, type of toxin and the toxin receptors involved in the process (Blazek et al., 2015; von Hoven et al., 2019). The rapid change of intracellular ion concentration upon plasma membrane insult leads to an activation of different signaling pathways. In some cases, mitogen activated kinases (MAPK) are activated after membrane insult, which initiate the production of interleukins (IL) such as IL-4, IL-6 or tumor necrosis factor α (TNFα) for host cell protection as well as signaling to cells of the immune system (Chow et al., 2010; Fedele et al., 2010). At the same time, activation of MAPK is connected to the initiation of death pathways such as apoptosis, whereby in human endothelial cells a cleavage of procaspases 6, and 9, and cleavage of procaspase 3 was detected, leading to DNA fragmentation (N’Guessan et al., 2005). Ca^2+^ influx also leads to the recruitment of peripheral lysosomes to the plasma membrane, which in turn causes exocytosis of membrane repair enzymes such as lysosomal ASM. This process is accompanied by short-term externalization of lysosomal-associated membrane protein 1 (LAMP1) (Reddy et al., 2001). Externalized ASM cleaves sphingomyelin in the outer leaflet of the plasma membrane thereby generating ceramide-enriched platforms (Grassmé et al., 2001a), which serve to cluster receptors (Grassmé et al., 2001b), but also cause an ATP-independent invagination of the plasma membrane (Babiychuk et al., 2008) and a subsequent endocytosis of the pore (Idone et al., 2008). Endocytosis of α-toxin pores had been identified previously to reseal the plasma membranes of cells intoxicated with staphylococcal α-toxin (Husmann et al., 2009) and thus represents one additional pathway of plasma membrane repair next to the removal of pores by vesicle budding and the clogging by annexins (reviewed, e.g., in Bischofberger et al., 2012; Brito et al., 2019).

We here show that treatment with staphylococcal α-toxin induces the recruitment of LAMP1 positive vesicles to the extracellular environment of endothelial cells, which thereby release the membrane repair enzyme ASM into the extracellular environment. This recruitment coincides with elevation of cytosolic calcium levels suggesting that α-toxin triggers membrane repair mechanisms in host cells as they were observed for other pore-forming toxins.

## Materials and Methods

### Staphylococcus aureus toxins

Recombinant *S. aureus* toxins were purchased from Gentaur GmbH (Aachen, Germany). α-toxin was purchased from Gentaur GmbH or Sigma (Taufkirchen, Germany).

### Bacterial culture

*S. aureus* strains (Table S1) were grown on trypticase soy agar (TSA) plates or TSA containing 5% defibrinated sheep blood. Appropriate antibiotics were added when selection was required. For infection experiments, *S. aureus* strains were grown over night in brain heart infusion (BHI) medium with appropriate antibiotics. Overnight cultures were incubated aerobically under constant shaking (600 rpm) at 37°C. For preparation of bacterial supernatants, overnight cultures were centrifuged for 2 min at 14,000 rpm and the supernatant was sterile-filtered through a syringe filter with a pore size of 0.2 µm.

### Mammalian cell cultures

All cell lines were grown mycoplasma-free in media supplemented with 10% fetal calf serum and penicillin/streptomycin (100 U/ml and 100 µg/ml, respectively). The murine endothelioma cell line bEnd.3 (ATCC® CRL-2299) was grown in Dulbecco’s Modified Eagle’s Medium (DMEM), supplemented with 2% L-Glutamine. Human monocytic THP-1 cells were seeded with a density of 2.5×10^5^ cells/well in a 24 well plate and challenged with sterile culture supernatants or purified alpha-toxin for 30 min at 37°C.

The human epithelial cells T24 (kindly provided by Prof. Alexandra Schubert-Unkmeir) and the human lung microvascular endothelial line HuLEC-5a (ATCC® CRL-3244TM) were grown in MCDB 131 Medium with a final concentration of 10% FBS and supplemented with 2.76 µM hydrocortisol, 0.01 ng/ml human epidermal growth factor (hEGF), 2 mM L-glutamine and microvascular growth supplement (25 ml of a 20x stock solution per 500 ml medium). All media were purchased from Thermo Scientific.

### Acid sphingomyelinase activity assays

#### Thin-layer chromatography-based ASM activity assay

ASM activity assays were based on a published protocol (Mühle and Kornhuber, 2017). Briefly, supernatants (SNT) of eukaryotic cells were collected in 1.5 ml centrifugation tubes and immediately stored on ice. The adherent cells were lysed with ASM lysis buffer (250 mM sodium acetate, pH 5.0, 0.1% Nonidet P-40, 1.3 mM EDTA, and 1x Roche Complete protease inhibitor cocktail) for at least 15 min and were scraped off with a pipette tip and collected. Protein concentrations were determined by Bicinchoninic Acid (BCA)-Assay (Pierce™ BCA Protein Assay Kit) according to the manufacturer’s instructions. ASM activity in the cell culture SNT was measured by addition of 100 µl SNT to 100 µl ASM SNT assay buffer (200 mM sodium acetate buffer pH 5.0, 500 mM NaCl, 0.2% Nonidet P-40, 500 µM ZnCl_2_). ASM activity in the whole cell lysate (LYS) was measured in 100 μl lysate volume, equivalent to 1 μg total protein, and addition of 100 µl ASM LYS assay buffer (200 mM sodium acetate buffer (pH 5.0), 500 mM NaCl, 0.2% Nonidet P-40). To each assay buffer solution, 58 pmol of BODIPY-FL-C_12_ Sphingomyelin (Invitrogen) was added (1:2000 dilution) and samples were either incubated for 24 h (supernatant samples) or for 4h (lysate samples) at 37°C under constant shaking (300 rpm). Assays were terminated by addition of 250 µl chloroform:methanol (2:1) and brief mixing of the samples. Samples were centrifuged (3 min, 13,000 *g*), 100 µl of the lower phase was collected and evaporated in a SpeedVac at 45°C for 20 min. Samples were resuspended in 10 µl chloroform:methanol (2:1) and were separated by thin-layer chromatography (TLC) on silica-gel plates (ALUGRAM® Xtra SIL G 5×10 cm). Separation was performed in sealed glass chambers containing chloroform:methanol (80:20). The fluorescence of lipids was detected with a Typhoon 9200 Scanner (Amersham) using excitation with the 532 nm Laser.

#### Radioactive ASM activity assay

After incubation of bEnd.3 cells with several different *S. aureus* toxins, ASM activity was determined using a radioactive assay. Cells were lysed in ASM lysis buffer II (250 mM sodium acetate, pH 5.0, 1% Nonidet P-40 and 100 µM ZnCl_2_) for 10 minutes on ice and then tip-sonicated for 10 seconds. Samples were diluted to 0.1 % Nonidet P-40 and incubated with [^14^C]sphingomyelin (0.05 mCi per sample; 52 mCi/mmol; Perkin Elmer) for 30 to 60 minutes at 37°C. Sphingomyelinase reaction was terminated by addition of chloroform:methanol (2:1, v/v). Phases were separated and the upper, [^14^C]choline chloride containing phase was collected and radioactivity was determined by liquid scintillation counting.

### Cytotoxicity detection

Mammalian cells were challenged with sterile bacterial supernatant at multiplicities of infection (MOI) or concentrations as indicated. Host cell supernatants were collected at indicated time points and centrifuged at 14,000 g for 2 min. For each sample, triplicates were transferred to 96 well cell culture plates and measurement of lactate dehydrogenase (LDH) release into growth medium (as direct indicator of cell membrane integrity loss) was performed with the Cytotoxicity Detection Kit^PLUS^ (Roche) according to the supplier’s manual.

### Hemolysis assays

Hemolysis of red blood cells (RBC) was performed with sheep and rabbit blood samples (Fiebig Nährstofftechnik, Germany). Briefly, 1 ml whole blood was centrifuged at 150 g for 5 min. Supernatant was carefully removed and cell pellet washed with 0.9% sodium chloride solution. This step was repeated two times. The washed cell pellet was then diluted to yield a 1% cell dilution. Washed, diluted blood was then incubated with diluted α-toxin or bacterial supernatants by shaking for 1h at 500 rpm and 37°C. Samples then were centrifuged for 1 min at 13.000 rpm to yield cell free supernatants. Triplicates of the sample supernatants were transferred to 96-well microplates and were analysed in a Tecan InfinitePro200 plate reader at 506 nm ± 9 nm. Relative hemolysis was calculated with reference to maximal lysis (0.1% Triton-X 100) and a negative control (0.9 % NaCl).

### Fluorescence microscopy techniques

HuLEC were seeded 2-7 days prior experiment in a density of 2×105 cells/ml on glass cover slips. After treatment, cells were fixed for 15 min on RT with 4% PFA and subsequently permeabilized with PBS/5% FBS/0.1% Triton X-100. Cells were then incubated with primary antibody (ZO-1 -Cell signaling #13663 1:400; VE-Cadherin – R&D MAB9381 1:500) overnight in a humid chamber. The following day, glass cover slips were rinsed thrice with PBS, and incubated with secondary antibody for 1h at RT. Cells were finally rinsed thrice with PBS and mounted on microscopy slides in Mowiol and dried overnight.

### Ca^2+^ measurement

HuLECs were seeded in a density of 3×10^4^ cells/well one day prior use in a µ-Slide 8 well live-cell chamber (Ibidi). The following day, cells were rinsed thrice in PBS and individually incubated with MCDB131 growth medium supplemented with 4 µM Fluo 3-AM (Invitrogen) for 25 min at 37°C. Cells were again rinsed with PBS and RPMI1640 without phenol red (ThermoFisher, #32404014) complemented with 10% FBS was added. Imaging was performed on a TCS SP5 confocal microscope (Leica Biosystems, Wetzlar, Germany) using a 63x oil-immersion UV objective with a numerical aperture of 1.4. Images were taken at 37°C with a resolution of 1024 x 1024 every 7.5 seconds for up to 25 minutes. Treatment with ionomycin, toxin or supernatants was performed during image acquisition after 10 images of untreated cells were recorded. Time series of the treated samples were analyzed with Fiji/ImageJ (Schindelin et al., 2012). Rectangular regions of interest (ROI) for each Fluo 3-AM labeled cell were defined. The average intensity over time was determined by the ImageJ plugin Time Series Analyzer (version 3). Fluorescence intensities were calculated relative to the initial time points (before addition of toxins, controls or bacterial supernatants) and were plotted using GraphPad Prism.

### LAMP1 recruitment

HuLECs were seeded in a density of 1×10^5^ cells/well four days prior the experiment. The cells were washed thrice with PBS and then challenged for 30 min at 37°C, 5% CO_2_. At the start of each challenge, LAMP1 monoclonal antibody (Santa Cruz, H4A3) in a concentration of 1 µg per 1×10^6^ cells was added. The cells were then rinsed five times with PBS and fixed by addition of 4% PFA. Cells were again rinsed thrice with PBS, permeabilized and blocked with PBS containing 0.1% saponin and 5% FBS for 1h at RT. Secondary antibody (Alexa Fluor 488-coupled anti-mouse IgG, Santa Cruz) was diluted 1:1000 in PBS/0.1% saponin/5% FBS/Hoechst 33258 and was left on samples for 1h at 37°C. Cells were finally washed five times with PBS, mounted in Mowiol and dried overnight at RT. Fluorescence was detected with a Leica TCS SP5 confocal laser scanning microscope (Leica Biosystems, Wetzlar, Germany) using a 63x oil-immersion objective (Numerical Aperture 1.4).

### Immunoblotting

bEnd.3 cells were seeded 48 h before the experiment in 24 well (1×10^5^ cells/well). Before α-toxin challenge, cells were washed thrice with HEPES buffered saline (H/S: 132 mM NaCl, 20 mM HEPES pH 7.3, 5.4 mM KCl, 1 mM CaCl_2_, 0.7 mM MgCl_2_, 0.8 mM MgSO_4_) or with calcium free phosphate buffered saline (PBS: 137 mM NaCl, 2.7 mM KCl, 1.15 mM KH_2_PO_4_, 6,5 mM Na_2_HPO_4_). After indicated times of toxin incubation, cells were lysed for 5 minutes in SDS lysis buffer (25mM HEPES, 0.1 % SDS, 0,5 % deoxycholate, 1 % triton, 10 mM EDTA, 10 mM NaPP, 10mM NaF, 125 mM NaCl, 10 mM NaVO_4_, 10 µg/ml aprotinin/leupeptin) on ice. Samples were centrifuged at 20,800 g for 5 minutes at 4°C and supernatants were added to 5x SDS Laemmli buffer and boiled. An equivalent of about 6,000 cells of each sample was separated by 10 % sodium dodecyl sulphate polyacrylamide gel electrophoresis (SDS-PAGE). Proteins were blotted onto nitrocellulose membranes at 4°C overnight. Membranes were blocked in Starting Block Tris-buffered saline (TBS) blocking buffer (Thermo-Fisher Scientific, Darmstadt, Germany) for 1h and incubated with antibodies specific for p38 (Cell Signaling Technology, #9212) or phospho-p38 (Cell Signaling Technology, #9211), respectively. All antibodies were diluted 1:1000 in blocking buffer and were incubated for 1h at room temperature. Blots were washed five times with TBS/0.05% Tween 20 (TBS/T), incubated with alkaline phosphatase coupled anti-rabbit antibody (Abcam, Berlin, Germany) for 1h, washed again five times with TBS/T and afterwards two times in alkaline wash buffer, and developed with CDP-STAR with Nitro-Block II Enhancer system (Perkin Elmer, Rodgau, Germany).

HuLEC were seeded with a density of 2×10^5^ cells/well in a 12 well-plate and cultivated for 3 days. Cells were washed thrice in PBS and treated with 1 µg/ml α-toxin, or 10% sterile-filtered *S. aureus* JE2 SNT for 15 min in cell culture medium (MCDB131 medium containing 2 mM L-glutamine and 10% FBS inactivated at 70 °C). After treatment, cells were washed once with PBS on ice and lysed by adding per well 80 µL 2x Laemmli buffer (100 mM Tris-HCL pH 6.8, 4% SDS, 20% glycerol, 2% beta-mercaptoethanol, 25 mM EDTA, 0.04% bromophenol blue.). Lysates were incubated for 10 min on 95°C and separated by 10% SDS-PAGE. Semi-dry Western transfer on a PVDF membrane (Sigma) was performed for 2h at RT with 60 mA/10V per blot. Membranes were blocked for 1h in 5% bovine serum albumin (BSA; Albumin Fraktion V, Roth) in TBS/T at room temperature. Incubation with primary antibody (p38 [CST #9212] and phospho-p38 [CST #9215], 1:500 dilution) was performed overnight at 4°C. Membranes were washed thrice in TBS/T and incubated with secondary horseradish peroxidase-conjugated antibody (Dianova 111-035-144, 1:2500 in 5% BSA in TBS/T) for 1h at room temperature. Subsequently, blots were washed thrice with TBS/T, developed using the ECL system (Pierce) and imaged on an Intas HR 16-3200 chemiluminescence reader.

### Statistics

One way analysis of variance (ANOVA) or Kruskal-Wallis rank sum tests determined significant differences in the analyzed data, followed by Student’s t-tests using Graphpad Prism.

## Results

### Staphylococcal α-toxin leads to p38 phosphorylation and alterations in cytosolic Ca^2+^ levels in endothelial cells

Murine endothelia have been shown to degrade tight junctions upon challenge with staphylococcal α-toxin thereby leading to lung edema in vivo (Seeger et al., 1990; Bartlett et al., 2008; Becker et al., 2018). We here treated bEnd.3 endothelial cells with staphylococcal α-toxin and investigated phosphorylation of p38 MAPK, which has been previously shown to be activated by α-toxin in epithelial cells (Eiffler et al., 2016). Ten to twenty minutes after addition of the toxin we observed an increase in phospho-p38 (Fig.S1 A). This increase lasted for about 60 minutes, after which phosphorylation of p38 decreased. Phosphorylation of p38 was abolished when host cells were treated with the ADAM10 inhibitor GI254023X (Selleck Chemicals, Biozol, Germany) prior to toxin addition (Fig. S1B).

We next investigated, if p38 phosphorylation depended on the extracellular divalent cations Ca^2+^ and Mg^2+^.We therefore added α-toxin to host cells in H/S buffer or Ca^2+^/Mg^2+^-free PBS. Cells incubated in medium without divalent cations showed no change in α-toxin-dependent p38 phosphorylation (Fig.S1C), demonstrating that the α-toxin-dependent increase in phospho-p38 at the 15 min time point required divalent cations in the medium.

We then tested α-toxin dependent activation of p38 in a human microvascular endothelial cell line (HuLEC) and observed an increase of relative phospho-p38 levels in HuLEC treated with 1 μg/ml purified α-toxin (1.81 ± 0.46 fold relative to the medium control), wild-type (WT) *S. aureus* culture supernatant (SNT; 2.12 ± 0.45 fold), and SNT of the Δ*hla* mutant complemented with 1 μg/ml α-toxin (2.21 ± 0.51). Δ*hla* SNT was leading to significantly less p38 activation (1.46 ± 0.31 fold relative to control; p=0.0068) when compared to WT SNT or purified α-toxin (p=0.033), whereas differences in normalized fold-changes between either purified α-toxin (p=0.064) or WT SNT (0.61) and complemented Δ*hla* SNT were not significant (n.s.; Fig. 1A). This demonstrated that α-toxin activated p38 MAPK in the HuLEC in a similar fashion as observed for the murine endothelial cell line bEnd.3 (Fig S1).

**Fig. 1:**
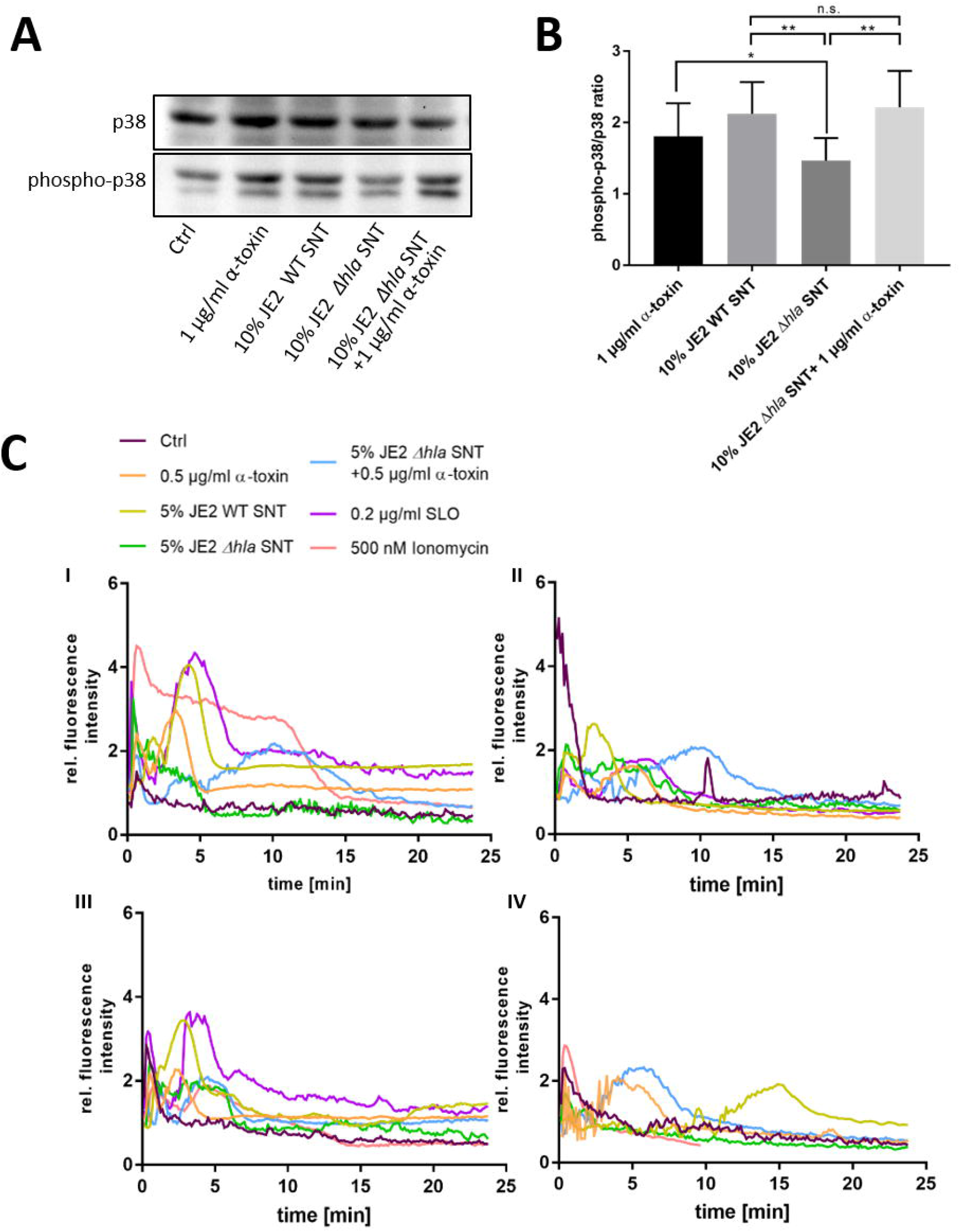
Human microvascular endothelial cells (HuLEC) treated with α-toxin induce phosphorylation of p38 MAPK. (**A, B**) HuLEC were treated with 1 µg/ml α-toxin, or SNT of *S. aureus* WT, a Δ*hla* mutant or Δ*hla* SNT complemented with 1 μg/ml α-toxin and phosphorylation of p38 was determined densitometrically by Western blot. Ratios of phospho-p38/p38 ratios were calculated and normalized to the untreated control, whose ratio was set to 1. (**C**) Calcium-influx is detected after α-toxin challenge. HuLEC cells were incubated with 4 µM Fluo 3-AM for 25 min at 37°C/5% CO_2_and subsequently analysed for calcium-specific increase of fluorescence by time-lapse imaging. After 75 sec (10 time frames), cells were treated with 500 nM Ionomycin, 0.2 μg/ml SLO (3.32 nM), 0.5 µg α-toxin (15 nM) or 5% supernatant of JE2 WT (JE2 SNT) or mutant JE2 lacking α-toxin (JE2 Δ*hla* SNT), or JE2 Δ*hla* SNT complemented with 0.5 μg/ml α-toxin. Shown are four independent experiments (I-IV).

Next, we investigated by time lapse imaging α-toxin-dependent changes in cytosolic Ca^2+^ levels in HuLEC (Fig. 1C, Supplemental movie 1). Cells reproducibly reacted to bacterial supernatants of wild-type *S. aureus* and purified α-toxin with an increase in intracellular Ca^2+^ levels (Fig. 1C, Supplemental movie 1). These elevations in cytosolic, Ca^2+^depended on the presence of α-toxin, although the time points of the Ca^2+^ peaks varied between the experiments. Treatment with bacterial supernatants lacking α-toxin (JE2 Δ*hla* SNT) prevented the strong Ca^2+^ signal in the endothelial cells, which was reinstated by adding 0.5 μg/ml α-toxin to culture SNT of the Δ*hla* strain. We observed a similar Ca^2+^ signal when Fluo-3 loaded cells were treated with SLO, another proteinaceous PFT (Fig. 1C). This SLO amount is equivalent to one fifth of the concentration of α-toxin monomers, demonstrating the differential sensitivity of HuLEC cells for both toxins.

For complementation of Δ*hla* SNT with purified α-toxin we determined the cytotoxic and hemolytic potential of bacterial supernatants and compared it with a commercially available preparation of α-toxin. Cytotoxicity was tested with the monocytic cell line THP-1. Cytotoxicity of bacterial SNT from the WT *S. aureus* strain JE2 was reduced in the Δ*hla* mutant (p=0.014) which was reflected by a similar increase in cytotoxicity upon addition of 1 μg/ml α-toxin to medium. Supernatants of the non-cytotoxic strain Cowan I (p=0.0001) and a JE2 mutant in the two-component system *sae*R (Δ*sae*R, p=0.0002) were drastically reduced (Fig. S2A). Our data thus suggests that the residual cytotoxicity observed for the Δ*hla* mutant is due to Sae-dependently produced leukocidins, which have been shown to target monocytic cells such as THP-1. Hemolysis was tested on washed blood from rabbit (Fig. S2B, C) and sheep (Fig. S2D). A titration of haemolytic activity of different toxin concentrations on rabbit RBC determined half-maximal hemolysis at 0.375 μg/ml for our preparation of the PFT. 0.5 μg/ml toxin of were approximately equivalent to a 5% dilution of WT SNT (Fig. S2C). Similar results were obtained for RBC derived from sheep blood. As expected rabbit RBC were more sensitive for α-toxin hemolysis when compared to sheep RBC (Fig. S2D).

### Staphylococcal α-toxin induces exocytosis of lysosomes

Since one Ca^2+^-dependent pathway of membrane repair after PFT insult involves exocytosis of peripheral lysosomes (reviewed in Bischofberger et al., 2012; Brito et al., 2019), we tested whether the supernatant of *S. aureus* cultures would transiently recruit lysosomal vesicles to the cell surface. In order to test this, we added antibodies directed against the lysosomal membrane protein LAMP-1 to the culture medium and left the cells untreated (Fig. 2A) or treated the cells with purified α-toxin (Fig. 2B), ionomycin (Fig. 2C), or bacterial supernatants of wild-type bacteria or an isogenic α-toxin mutant. After 30 minutes, we fixed and permeabilized the cells, and detected anti-LAMP-1 antibodies with fluorescently labeled secondary antibodies. We thereby specifically detect LAMP-1 proteins that were surface-exposed during the course of the challenge (Fig. 2D). By fluorescence microscopy we enumerated LAMP1 signals in the cells and observed significantly increased numbers of LAMP-1 positive vesicles in cells treated with ionomycin and supernatants from α-toxin-proficient bacteria when compared to supernatant from the Δ*hla* mutant (Fig. 2E).

**Fig. 2:**
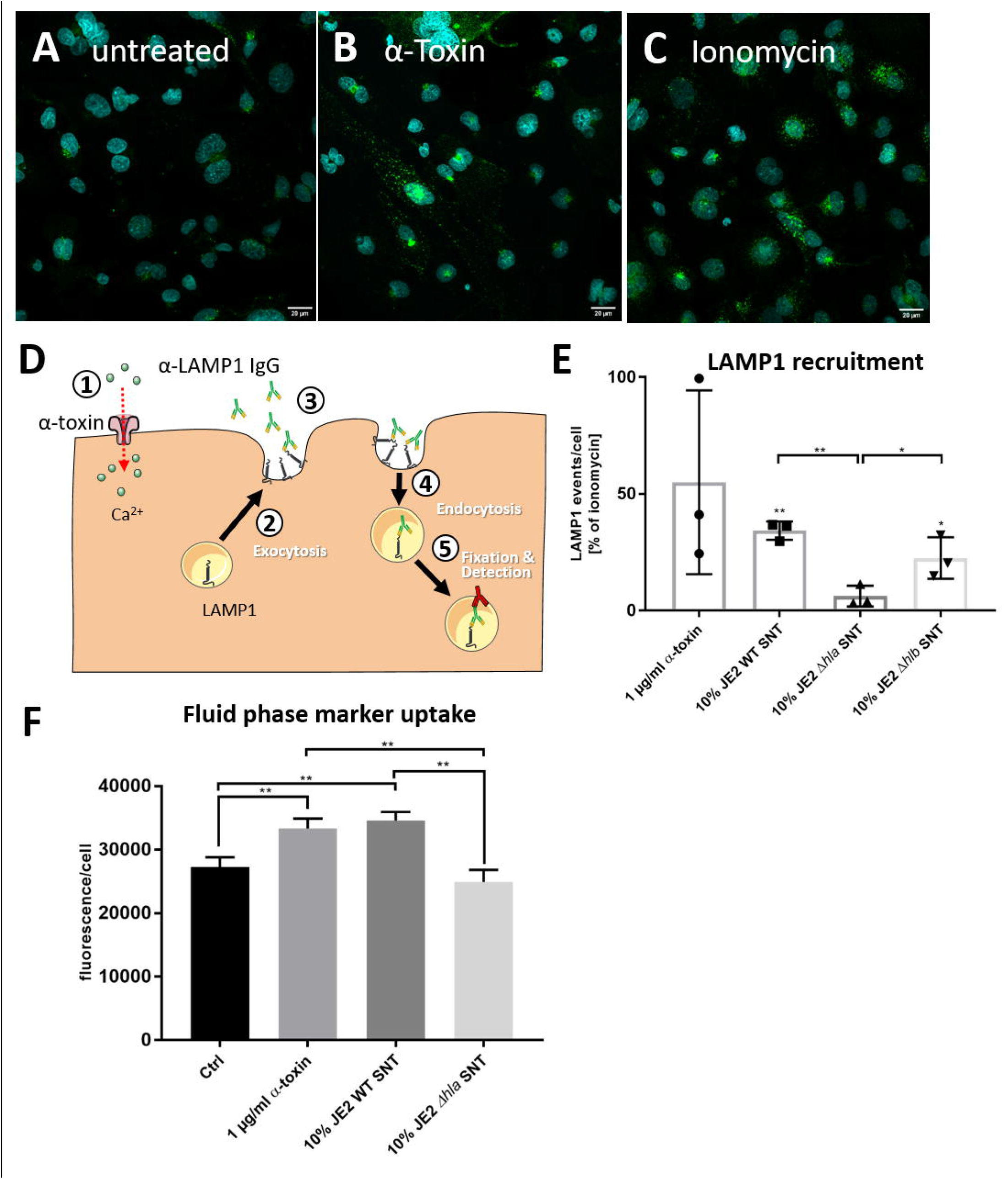
*S. aureus* induces recruitment of LAMP-1 positive vesicles to the plasma membrane. HuLEC cells were left untreated (**A**) or were treated for 30 min with 1 µg/ml purified α-toxin (**B**) or 1 µM of the Ca^2+^ ionophore ionomycin (**C)**. Fluorescence microscopy in untreated cells and cells treated with α-toxin or ionomycin demonstrates more and stronger LAMP1 signals in the treated cells. (**D**) Assay schematic: 1 µg LAMP1 antibody per 1×10^6^ cells was added together with α-toxin ①. Either by directly establishing a Ca^2+^ permissive pore or indirectly by activating endogenous host Ca^2+^ channels, the cytosolic Ca^2+^ concentration rises. Thereby, the antibodies can recognized exocytosed lysosomal membranes ② currently displayed at the host cell surface ③ as well as LAMP1-molecules which were taken up again during the course of the treatment ④. After fixation and permeabilization, an Alexa488-labelled secondary antibody was added and LAMP1 was detected by fluorescence microscopy ⑤. (**E**) Flow cytometric quantification of LAMP1-positive vesicles left untreated, or treated with α-toxin-and ionomycin, or cells treated with 10% *S. aureus* culture supernatants (SNT) of wild-type bacteria (JE2) or its isogenic α-toxin mutant (JE2 Δ*hla*) demonstrates that the LAMP1 surface recruitment is abrogated in the Δ*hla* strain. (**F**) Epithelial cells increase uptake of a membrane impermeable fluid phase dye after challenge with α-toxin. T24 cells were incubated in cell culture medium containing Alexa633-hydrazide and either 1 µg/ml purified *S. aureus* α-toxin or 10% SNT from JE2 WT or its isogenic Δ*hla* mutant was added. After 30 min cells were detached by trypsinization and fluorescence was analysed by flow cytometry. For statistical analysis, experiments in E and F were independently conducted three times (n=3). A Kruskal-Wallis rank sum test determined significant differences in the analyzed data, which were further tested with pairwise Student’s t-tests. *: p<0.05, ** p< 0.01; ***: p<0.001. For more details see text.

We next added the membrane impermeable fluorescent dye Alexa633 hydrazide to the cell culture medium. The dye thus served as fluid phase marker. We then added either 1 µg/ml purified *S. aureus* α-toxin or 10% bacterial culture supernatant from either *S. aureus* JE2 wild-type (WT) or its isogenic Δ*hla* mutant for 15 min. After the incubation, cells were washed, detached by trypsinization and analysed for Alexa633 fluorescence by flow cytometry. We observed an increased uptake of the membrane impermeable fluid phase dye after challenge with purified α-toxin or *S. aureu*s WT SNT, whereas fluorescence of cells treated with SNT from the Δ*hla* mutant were at the level of the negative control (Fig. 2E).

Thus our data suggest that upon treatment with α-toxin or bacterial SNT containing the toxin, lysosomes are transiently recruited to the cell surface.

### ASM is released into the extracellular environment by human endothelial cells upon challenge with α-toxin

We have previously shown that the acid sphingomyelinase is activated rapidly after staphylococcal α-toxin challenge in macrophages and endothelial cells (Becker et al., 2018). Therefore, we investigated ASM activity in host cells by treatment with sterile bacterial supernatants. For that purpose we adapted a previously published ASM assay (Mühle and Kornhuber, 2017) which is based on separation of fluorescently labeled C12 sphingomyelin and ceramide by thin-layer chromatography. However, we observed a prominent ASM activity in our cell culture medium, which was caused by fetal bovine serum (FBS) (Fig. S3). We here used culture supernatants of *S. aureus*, which are known to contain copious amounts of short amphiphilic cytolytic peptides, the so called phenol-soluble modulins (PSM) (Surewaard et al., 2012). We therefore had to include FBS in the cell culture media, since serum lipoproteins inhibit host cytolysis by PSM. Heat-inactivation of FBS at 55°C for 30 min (Spence et al., 1989) reduced ASM activity, but only heat-treatment of FBS for 1 hour at 70°C resulted in complete loss of ASM activity, whereas PSM-dependent cytolysis was still abrogated under these conditions (Fig. S4).

Using the activity assays described above, we identified an additional background sphingomyelinase activity. The *S. aureus* genome harbors the structural gene for a bacterial sphingomyelinase (bSMase) β-toxin (β-hemolysin, *hlb*). We found sphingomyelinase activity not only in culture supernatants from the strongly β-toxigenic strain 6850, but also in the MRSA strain JE2, suggesting that the latter strain had an excised bacteriophage. bSMase activity was only completely abolished in JE2 mutants, which carried an additional transposon insertion within the structural *hlb* gene. By contrast, the bacterial phospholipase C (*plc*) activity did not contribute to sphingomyelin degradation (Fig. S5). Compared to supernatants from *S. aureus* strains 6850 and RN4220, sphingomyelinase activity of the clinically relevant *S. aureus* strain JE2 was smaller but still detectable.

With the adapted assay conditions we next compared effects of bacterial culture supernatants from *S. aureus* or purified toxin on endothelial cells and challenged the cells with 10% bacterial culture supernatant of wild-type *S. aureus* JE2, or its isogenic Δ*hla* or Δ*hlb* mutants, or treated the cells with 1 µg/ml purified α-toxin. At different time points during a 30-minute incubation we collected both cell culture medium and cell lysates. Both fractions were assayed for ASM activity (Figure 3A-D). We thereby observed an α-toxin-dependent increase in host ASM activity in the culture medium over time, whereas ASM activity in cell lysates decreased (Fig. 3A-D). The absence of lactate dehydrogenase (LDH) in the media indicated that the endothelial cell membranes remained intact during the treatment (Fig. 3E) suggesting that ASM is released by peripheral lysosomes that fuse with the plasma membrane. ASM release by HuLEC was also induced by treatment with streptolysin O (SLO) as well as the calcium ionophore ionomycin (Fig. 3F), which again is reminiscent of membrane repair processes leading to ASM release (Andrews et al., 2014). The residual sphingomyelinase activity observed in SNT of the Δ*hla* strain may be derived from the bacterial SMase β-toxin, which can be expressed by *S. aureus* (Fig. S5, see also above). We also tested an *hlb* insertional mutant of *S. aureus* JE2, which was β-toxin deficient (Fig. S5). Challenging endothelial cells with supernatant of this mutant strain showed similar dynamics of ASM activation when compared to cells challenged with supernatants of the wild-type strain. This demonstrated that the bSMase does not contribute to the observed effects (Fig. 3C).

**Fig. 3:**
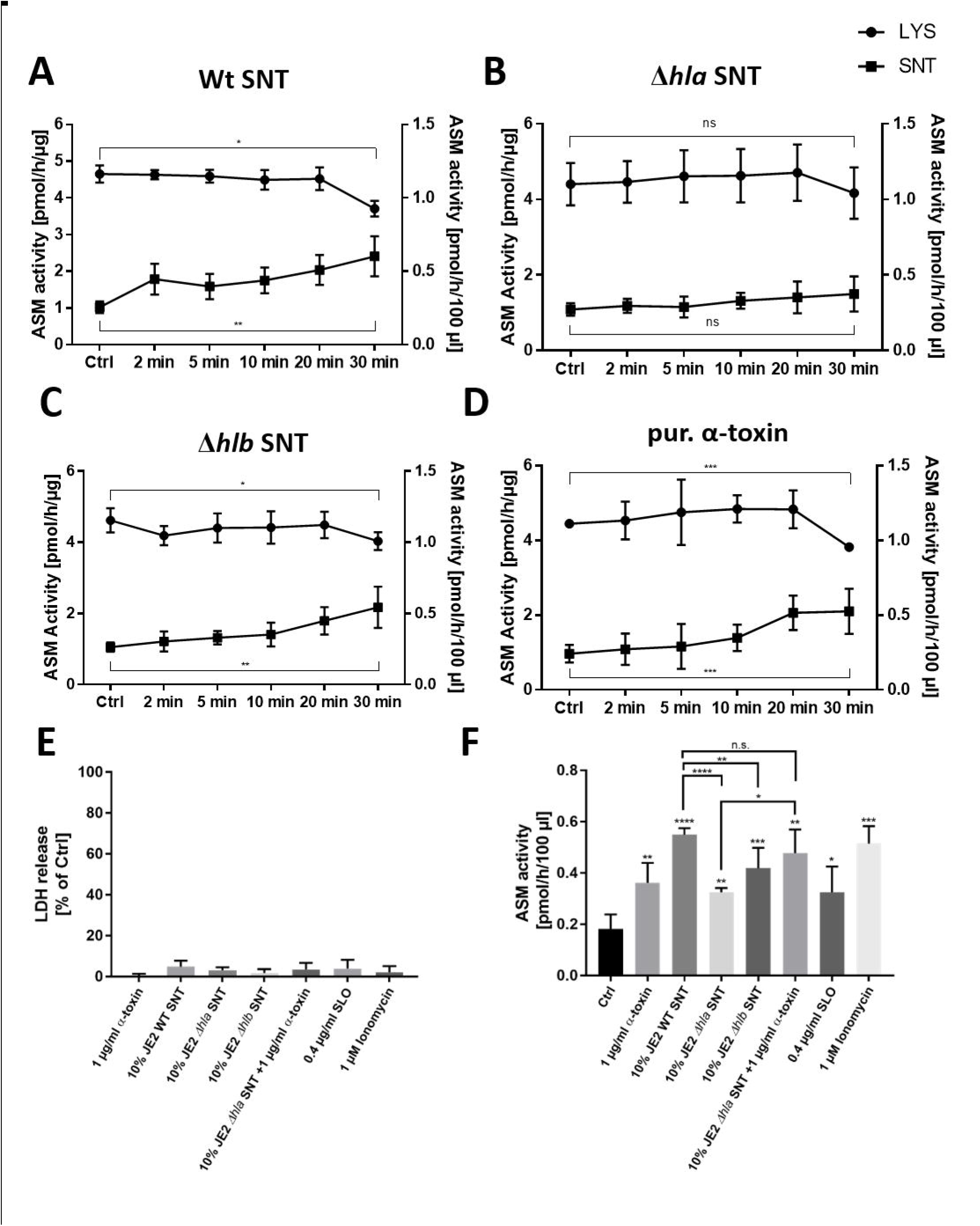
Staphylococcal α-toxin induces ASM release into the supernatant of endothelial cells: HuLEC were either challenged with 10% bacterial culture supernatant of (**A**) *S. aureus* JE2 WT (SNT WT), (**B**) JE2 lacking α-toxin (Δ*hla* SNT) or (**C**) JE2 lacking β-toxin (Δ*hlb* SNT), or were treated with 1 µg/ml of purified α-toxin (**D**) or were left untreated (Ctrl). ASM activity was measured at the indicated time points in cell culture medium (“SNT”, closed squares) as well as in endothelial cell lysate (“LYS”, closed circles). ASM activity of lysates are given in pmol/h/μg (left y-axis) and were normalized to the total protein amount in cell lysates. ASM activities in culture media are given as pmol/h/100 μl (right y-axis). **E**) LDH assays of the culture supernatants in **A-D** demonstrate that the endothelial cells remained intact over the course of the experiment. **F**) HuLEC were challenged for 30 min with different samples. The reduced ASM release in the Δ*hla* SNT is complemented by external addition of 1 g/ml α-toxin and ASM release is also induced by 0.4 μg/ml SLO and 1 μM ionomycin. **E, F**) n=5 for Ctrl, 1 μg/ml α-toxin, WT SNT and Δ*hlb* SNT; n=3 for the remaining samples. Significance was calculated with Student’s t-test. *: p<0.05, ** p< 0.01; ***: p<0.001; ns: not significant.

Thus, our results demonstrate that peripheral lysosomes are recruited to the endothelial cell surface upon addition of staphylococcal α-toxin suggesting induction of a membrane repair mechanism that involves the release of lysosomal ASM. In line with this, the challenge of endothelial cells with supernatants lacking α-toxin failed to recruit ASM to the extracellular space. This indicated that α-toxin is the major toxin involved in this process in endothelial cells. Supporting this hypothesis, we observed that HuLEC cells, which were grown for seven days on a collagen matrix and displayed intact tight junctions, were not lysed by bacterial culture supernatants of the *S. aureus* JE2 Δ*hla* mutant, whereas supernatant collected from the wild type readily lead to tight junction breakdown and endothelial cell lysis (Fig. S6). Since our previous study determined that α-toxin induced ASM activity in murine bEnd.3 cells, we also tested, if other staphylococcal pore-forming toxins would lead to ASM activation in bEnd.3 cells. We thus treated the cells with combinations of purified subunits of the bi-component pore-forming toxins hemolysin γ (HlgA+HlgB, or HlgC+HlgB, respectively) or the leukocidins LukDE or LukSF (Fig. S7). None of these combinations led to ASM activation over the course of a 60-minute treatment. This again demonstrates that α-toxin is the main staphylococcal toxin leading to ASM release by endothelial cells.

## Discussion

α-toxin is secreted by *S. aureus* as a monomer and is assembled as a heptameric protein complex after binding to non-specific lipid receptors (Schwiering et al., 2013; Ziesemer et al., 2019) or the metalloprotease ADAM10 (Wilke and Bubeck Wardenburg, 2010) on target membranes. Due to its small pore diameter, it was suggested that *S. aureus* α-toxin pores are only permeable for small cations, predominantly potassium (Harshman et al., 1989; Jonas et al., 1994), although the permeability of the plasma membrane for Ca^2+^ and larger molecules such as ATP was also demonstrated (Jonas et al., 1994; Gierok et al., 2014). A hallmark consequence of cell intoxication with pore-forming toxins is the uncontrolled exchange of ions between the extra-and intracellular space (Dal Peraro and van der Goot, 2016). Intracellular concentration changes of specific ions induce tailored cell responses in order to re-establish homeostasis (Cooper and McNeil, 2015). When α-toxin elicits Ca^2+^ influx, activation of kinases critical for cell autonomous defense mechanisms ensues (Rosen et al., 1994; Liu et al., 2016). Central to this mechanism is the phosphorylation of the mitogen-activated protein kinase p38 (p38 MAPK), which is activated upon exposure of mammalian cells to the staphylococcal toxin (Husmann et al., 2006; Kloft et al., 2009; Eiffler et al., 2016) and which is also important for target cell recovery from the toxin insult (Husmann et al., 2006). Accordingly, we detected phosphorylation of p38 MAPK after incubation with purified staphylococcal α-toxin in human (Fig. 1) and murine endothelial cells (Fig. S1). p38 phosphorylation was reduced upon removal of extracellular divalentions or upon inhibition of the metalloprotease ADAM10 (Fig. S1), the receptor for α-toxin on target membranes (Wilke and Bubeck Wardenburg, 2010). Further, it has been demonstrated by others that α-toxin heptamerization is inhibited upon addition of the ADAM10 antagonist GI254023X (Inoshima et al., 2011), whereas the protease activity of ADAM10 is not required for α-toxin pore formation (Ezekwe et al., 2016). We here observed that endothelial cell p38 phosphorylation in presence of α-toxin was abrogated by pharmacological inhibition of ADAM10 with GI254023X (Fig. S1B).

Upon α-toxin treatment we observed a rise in intracellular Ca^2+^ concentrations, which peaked in presence of purified α-toxin or bacterial SNT containing the PFT (Fig. 1C; Supplemental Movie 1). We observed similar peaks of Fluo-3 fluorescence when applying the PFT streptolysin O (Fig. 1C). By contrast, treatment with the Ca^2+^ ionophore ionomycin led to a quickly appearing and strong Ca^2+^ peak (Fig. 1C; Supplemental Movie 1). This strong Ca^2+^ influx after ionomycin treatment has been detected over a variety of cell types (Yoshida and Plant, 1992; Mason and Grinstein, 1993; Gutierrez et al., 1999; Vasilev et al., 2012), whereas oscillating Ca^2+^ fluxes are often observed after mechanical cell injury (Reddy et al., 2001) and eventually result in calcium spikes (Justet et al., 2019). Thus our data suggests that α-toxin treatment of human and murine endothelial cells leads to p38 activation, which depends on divalent cations in the medium and and ADAM10It is currently unclear, if α-toxin itself thereby forms a Calcium-conductive channel or if challenge with the toxin results in activation of endogenous host Ca2+ channels, endogenous pores or even ruptures as has been discussed previously (von Hoven et al. 2019).

It has been observed that phosphorylation of p38 MAPK already occurs at low doses of α-toxin, whereas ADAM10 dependent breakdown of E-cadherin in toxin-challenged keratinocytes required higher concentrations of α-toxin (von Hoven et al., 2016). This may correspond with alternate states of membrane permeability, which might be caused by different α-toxin concentrations or the different cell types (von Hoven et al., 2019). This is further supported by the induction of Ca^2+^-independent membrane repair after α-toxin insult of fibroblasts (Valeva et al., 2000).

Plasma membrane disruption e.g. by PFT results in changes in intracellular ion-concentrations, which in turn trigger cell responses leading to membrane repair or cell death (reviewed in Bischofberger et al., 2012; Brito et al., 2019). p38 is implicated in survival pathways such as autophagy as well as plasma membrane repair after challenge with staphylococcal α-toxin (Husmann et al., 2006; Kloft et al., 2009; Kao et al., 2011) and interference with p38 MAPK-dependent membrane recovery pathways caused host cell death (Husmann et al., 2006). Calcium-dependent membrane repair pathways for instance include recruitment of annexins to damaged membrane areas where they assemble to clog pores. The survival mechanism against mechanical membrane injury (Reddy et al., 2001) as well as insult by some PFT involves the Ca^2+^-dependent exocytosis of peripheral lysosomes to the cell surface thereby releasing membrane repair enzymes such as lysosomal ASM into the extracellular environment. ASM-dependent conversion of outer leaflet sphingomyelin to ceramide facilitates endocytosis of the toxin pores (reviewed in Andrews, 2019; e.g. Brito et al., 2019).

In line with the latter repair strategy, our ASM activity assays revealed that intoxication with *S. aureus* α-toxin leads to a re-partitioning of ASM to the extracellular space (Fig. 3). These findings are supported by the increased exposure of the lysosomal membrane protein LAMP-1 on the plasma membrane after α-toxin challenge. This suggested that the ASM activity present in the cell culture media originated from peripheral lysosomes that fuse with the plasma membrane to deliver membrane repair enzymes.

LAMP1 exposure to the extracellular environment is described to be rather short, since these externalized lysosomal membranes are quickly re-endocytosed (Andrews, 2017). Accordingly, we observed increased rates of endocytosis of a fluorescent fluid phase marker, which was α-toxin-dependent in our observations (Fig. 2E). Since host cells also have been shown to recover from α-toxin injury by endocytosis of the affected membrane patches (Husmann et al., 2009), which is also observed for other PFTs (Idone et al., 2008), host cell plasma membrane permeabilized by staphylococcal α-toxin may dispose of the pores by recruitment of peripheral lysosomes, the accompanied release of repair enzymes such as ASM, as well as re-endocytosis of the additional membrane fractions.

ASM is a crucial factor for the conversion of sphingomyelin to ceramide in the extracellular leaflet of the injured plasma membrane (Andrews et al., 2014). Since this removal of plasma membrane sphingomyelin mediates resistance to α-toxin (Ziesemer et al., 2019) it is tempting to speculate that ASM release of injured mammalian cells do not only initiate plasma membrane repair mechanisms, but also may decrease membrane sensitivity against the toxin. In addition, α-toxin pores were found to be excreted, if cells were unable to remove the pore by proteolysis (Husmann et al., 2009). Thus, the processes of pore endocytosis, excretion as well as release of ASM to the culture medium may be interlinked in certain cell types.

In order to assess ASM activity in the culture supernatants of endothelial cells treated with staphylococcal culture supernatants, we had to include fetal bovine serum in order to inactivate phenol-soluble modulins, but were confronted with a strong background ASM activity in the serum. This activity was reliably inactivated by treatment at 70°C for 1 h (Fig. S3), while the serum was still inhibiting cytolysis by PSMs (Fig. S4). Further, we found that the commonly used laboratory strain RN4220 and the strongly cytotoxic strain 6850 could not be used for our ASM analyses due to their strong production of bacterial SMase. Although β-toxin is a neutral SMase (Doery et al., 1963), its activity is still detectable under acidic assay conditions. Clinical strains are often positive for so-called β-converting phages, which inactivate the bSMase by lysogeny (Goerke et al., 2006; Katayama et al., 2013). However, reactivation of the phage results in β-toxin expression (Salgado-Pabon et al., 2014; Tran et al., 2019) which was recently corroborated by bacterial single cell RNAseq in *S. aureus* culture (Blattman et al., 2020). We found sphingomyelinase activity in *S. aureus* JE2 was abrogated only by an additional insertional mutation in the *hlb* gene. Although bSMase activity in strain JE2 was rather low and only measurable in concentrated bacterial supernatants, we recommend testing each genetic background of *S. aureus* for a potential confounding SMase activity.

In summary, we here show – to our knowledge for the first time – that treatment with *S. aureus* α-toxin recruits acid sphingomyelinase to the extracellular space of endothelial cells. Our data further suggest that α-toxin is the major staphylococcal toxin causing the observed effects in these cells, since supernatant of a *S. aureus* α-toxin mutant did not induce ASM release and tight-junction disruption in endothelial cells. In addition, murine endothelial cells treated with other purified staphylococcal toxins did not show increase in ASM activity.

## Acknowledgements

We thank the Deutsche Forschungsgemeinschaft for funding this project within the research group FOR2123 (to E.G. and M.J.F.) and the training group GRK2581 (to M.J.F.). We further are indebted to Simon Peters and Alexandra Schubert-Unkmeir (Institute for Hygiene and Medical Microbiology, Universitätsklinikum Würzburg) as well as Kathrin Stelzner and Nadine Vollmuth (Chair of Microbiology, University Würzburg) for materials or valuable help with techniques.

**Table 1:** Strains used in this study.

| *S. aureus* Strain | Remarks | Source/Publication |
| --- | --- | --- |
| **JE2** | MRSA, USA300 | (Fey et al., 2013) |
| **JE2 $\Delta hla$** | JE2 with a markerless deletion of α-toxin | (Becker et al., 2018) |
| **JE2 *hly*::Tn** | Transposon-insertion mutant NE1261 re-transduced in JE2 wild-type genetic background | (Fey et al., 2013) |
| **JE2 *plc*::Tn** | Transposon-insertion mutant NE678 re-transduced in JE2 wild-type genetic background | (Fey et al., 2013) |
| **Cowan I** | Over-expression of protein A, Isolated by Cowan from a patient with septic arthritis in 1935; strongly cell invasive, yet non-haemolytic and non-cytotoxic | ATCC 12598 |
| **RN4220** | phage-cured, strong β-toxin producer, cloning strain | (Kreiwirth et al., 1983) |
| **6850** | strong α-toxin and β-toxin producer, highly cytolytic | (Vann and Proctor, 1987) |
| **6850 *hly*** | insertional mutant of *hly* in 6850 (CG33) | (Grosz et al., 2014) |

## Supplemental Figure legends

**Fig. S1: α-toxin induces p38 MAPK phosphorylation which is dependent on divalent cations in the medium and ADMA10:** bEnd.3 were incubated with 10 µg/ml α-toxin for 5-120 min and phosphorylation of p38 was determined by Western blot (**A-C**). Pre-treatment of cells with the ADAM10 inhibitor GI254023X inhibited p38 phosphorylation (**B**). Ca^2+^ is required for p38 activation, since a buffer lacking Ca^2+^ reduced α-toxin induced p38 phosphorylation. Shown are representative Western blots of three independent experiments. Phosphorylation of p38 was determined densitometrically and was normalized by signals measured for total p38. p38 is activated significantly during a period ranging from 10 through 60 min post intoxication (**A**). (**B**) Pre-treatment of cells with the ADAM10 inhibitor GI254023X inhibited p38 phosphorylation significantly at all investigated time points. (**C**) Ca^2+^ is required for p38 activation, which is observed at 15 and 30 min after intoxication. Activation is not observed in buffers lacking Ca^2+^. Statistical analysis was performed with GraphPad Prism using pairwise Student’s t-tests. *: p< 0.05; **: p< 0.001; ****: p< 0.0001.

**Fig. S2: THP-1 monocyte cytotoxicity and hemolysis of bacterial culture supernatants and α-toxin**. (**A**) Cytotoxicity of *S. aureus* JE2 SNT is reduced when compared to SNT of its isogenic α-toxin mutant (Δ*hla*), Cowan I, or a *sae*R mutant (Δ*sae*R), whereas 1 μg/ml α-toxin lysed host cells. Statistical analysis was performed with GraphPad Prism using One-way ANOVA and unpaired Student’s t-tests. *: p< 0.05; **: p< 0.001; ***: p< 0.0001. Hemolysis assays were performed using washed rabbit (**B, C**) and sheep blood (**D**). For details see main manuscript text.

**Fig. S3: FBS exhibits a strong ASM activity which is inactivated upon treatment at elevated temperatures**. (**A**) MCDB131 medium supplemented with untreated of heat inactivated (55°C for 30 min) FBS was tested for its ASM activity. Heat-inactivation for 30 min at 55°C reduced ASM activity. (**B**) Heat inactivation for 1h at 70°C (Medium 70°C) abolishes ASM activity of FBS completely. Thin-layer results are representative images of multiple experiments.

**Fig. S4: *S. aureus* phenol soluble modulins are inhibited by heat-treated FBS**. Epithelial cells were challenged with increasing concentrations of culture supernatants of *S. aureus* overnight cultures diluted in cell culture media. Additionally 70°C-treated FBS was added or the assays were left FBS-free. The heat-treated FBS shows the same inhibition of cytotoxicity as normal FBS. n=3. Significance was calculated with Student’s t-test. ***: p<0.001; ns: not significant.

**Fig. S5: *S. aureus* exhibits toxin specific sphingomyelinase activity**. Equal amounts of supernatants (SNT) from *S. aureus* overnight cultures were tested for sphingomyelinase activity. *S. aureus* exhibited a strain-specific conversion of sphingomyelin to ceramide with strains 6850 and RN4220 showing strongest bSMase activities. Even the clinical strain JE2 and demonstrates measurable activities, although the responsible ORF *hlb* is inactivated by phage lysogeny in this strain suggesting phage reactivation. Sphingomyelinase activity is absent from insertional mutants within *hlb* (termed “Δhlb”) in strain JE2 and 6850, as well as in strain Cowan I. Thin-layer results are representative images of three independent experiments.

**Fig. S6: *S. aureus* JE2 Δ*hla* supernatant does not induce tight junction degradation and endothelial cell death**. HuLEC were seeded 7 days prior to toxin challenge on glass cover slips coated with collagen. Cells were either treated with 10% bacterial culture supernatants of wild-type *S. aureus* JE2 (**A, B**) or its isogenic Δ*hla* mutant (**C, D**). Phase contrast microscopy images (**A, C**) were taken at 10x magnification. Immunofluorescence images (**B, D**) taken at 40x magnification. Blue – Hoechst 33258; Red – ZO-1; Yellow – VE-Cadherin.

**Fig. S7: Purified *S. aureus* toxins other than α-toxin fail to induce ASM activation in murine endothelial cells**. bEnd.3 cells were challenged with various combinations of hemolysin γ (Hlg) or leukocidin (Luk) subunits for 5 – 60 min and ASM activity was measured in cell lysate. n=3.

## Supplemental movie

**Supplemental video 1: Eleveated Calcium levels in host cytosol are α-toxin-dependent**. Human lung microvascular endothelial cells were incubated with the calcium-sensor 4 µM Fluo 3-AM for 25 min at 37°C/5% CO_2_ and subsequently analysed imaged every 7.5 seconds. Where indicated cells were treated with 500 nM ionomycin, 0.4 μg/ml SLO, 0.5 µg α-toxin or 5% of bacterial culture supernatants (SNT) of wild-type *S. aureus* JE2, its isogenic α-toxin mutant JE2 Δ*hla*, and JE2 Δ*hla* SNT complemented with 0.5 µg/ml α-toxin.

## Notes

### Competing Interest Statement

The authors have declared no competing interest.

