## Supplementary figures and images for "*Staphylococcus aureus* α-toxin induces acid sphingomyelinase release from a human endothelial cell line"

### Supplemental Figure 1

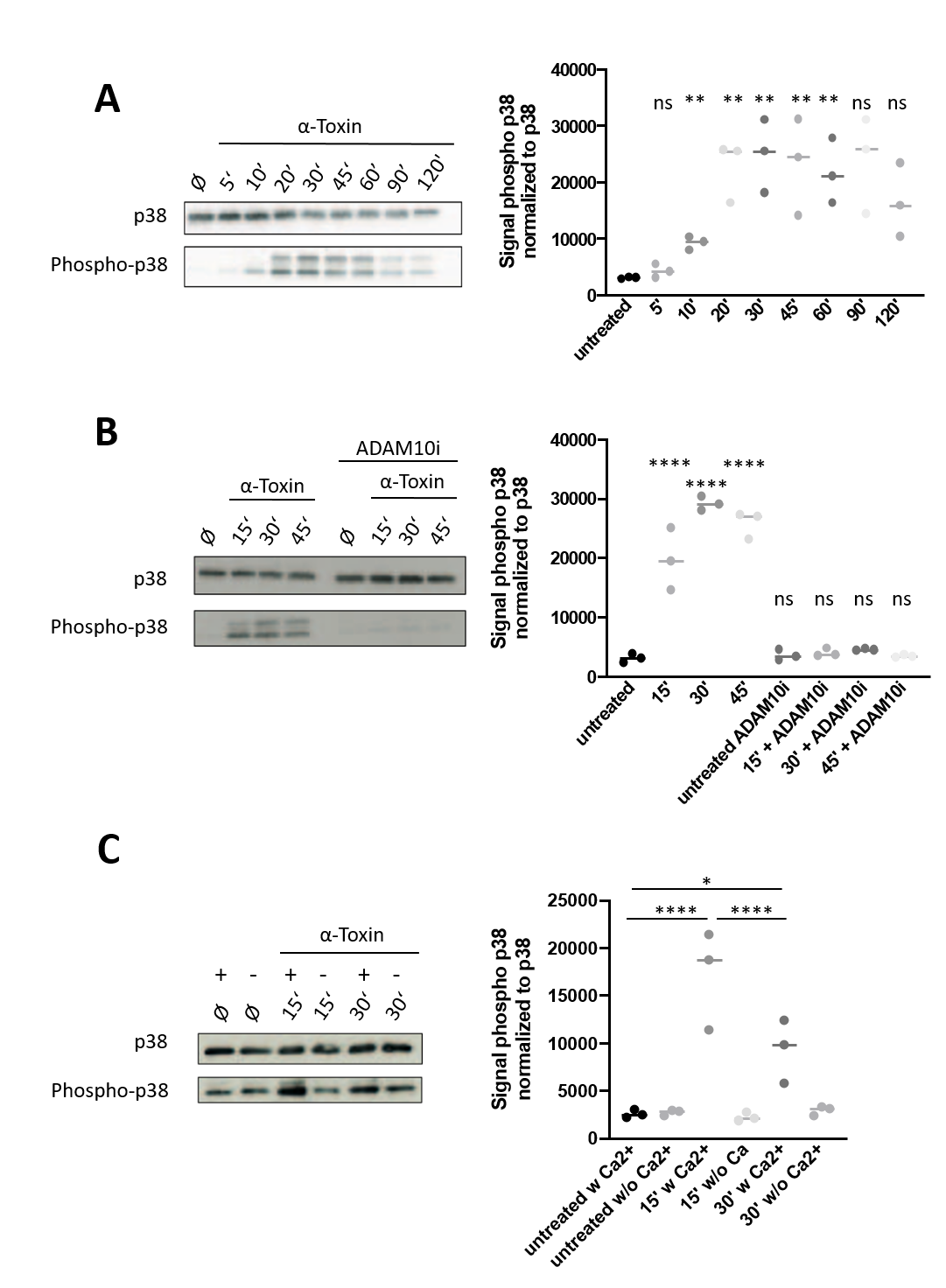

### Supplemental Figure 2

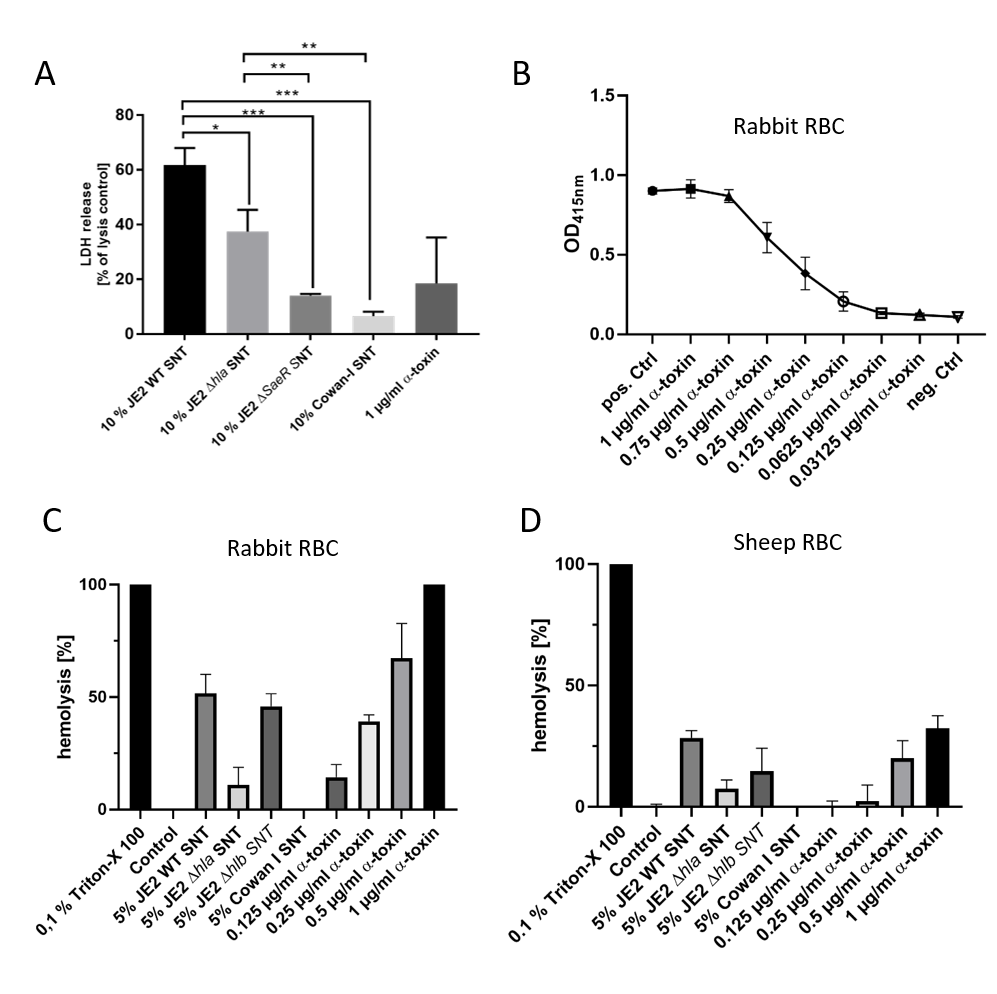

### Supplemental Figure 3

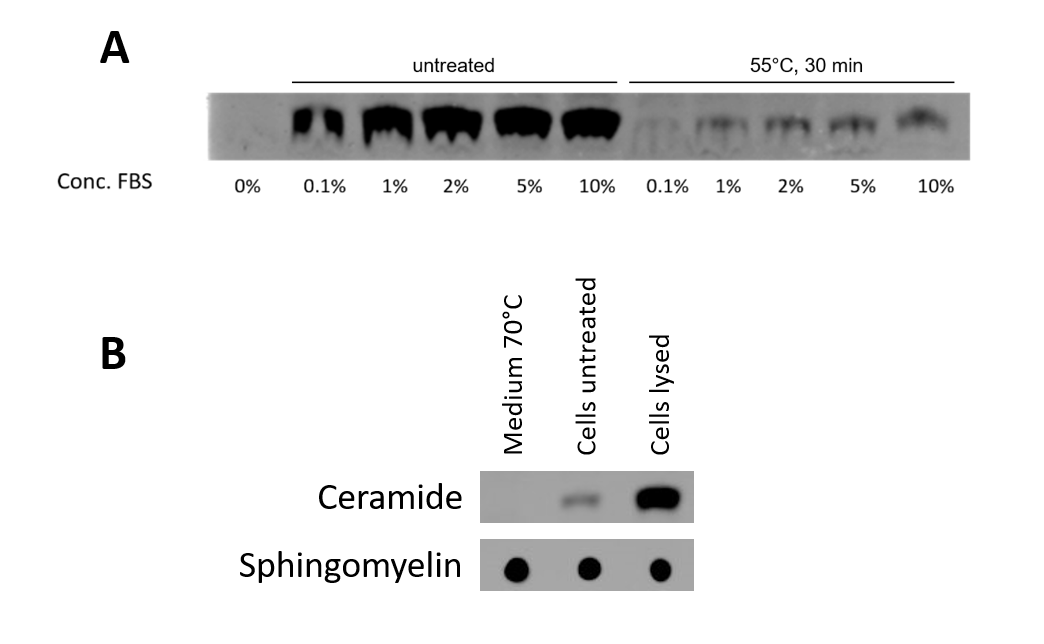

### Supplemental Figure 4

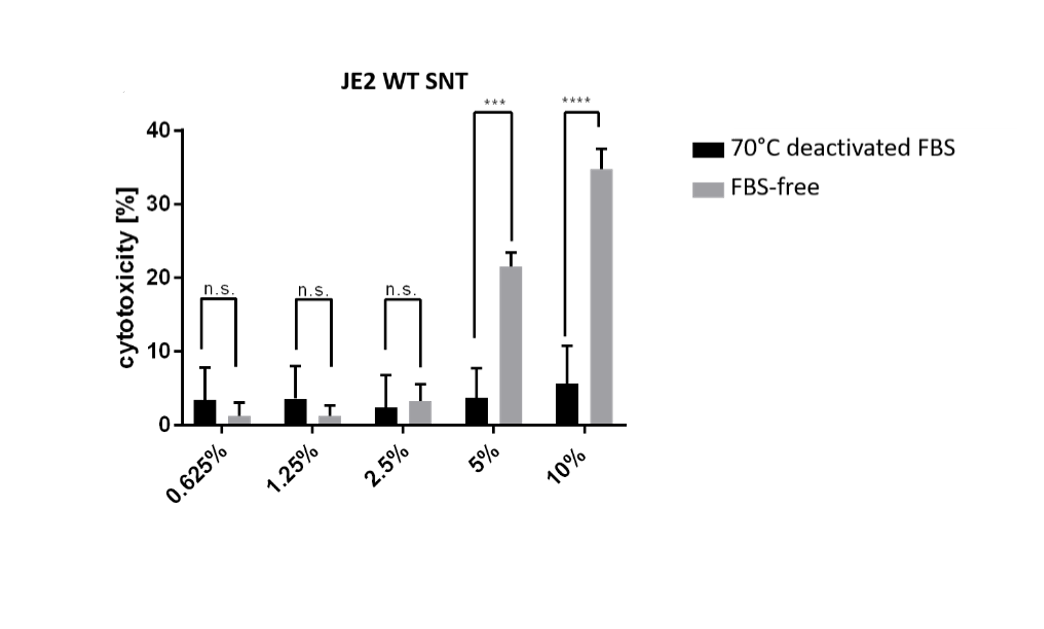

### Supplemental Figure 5

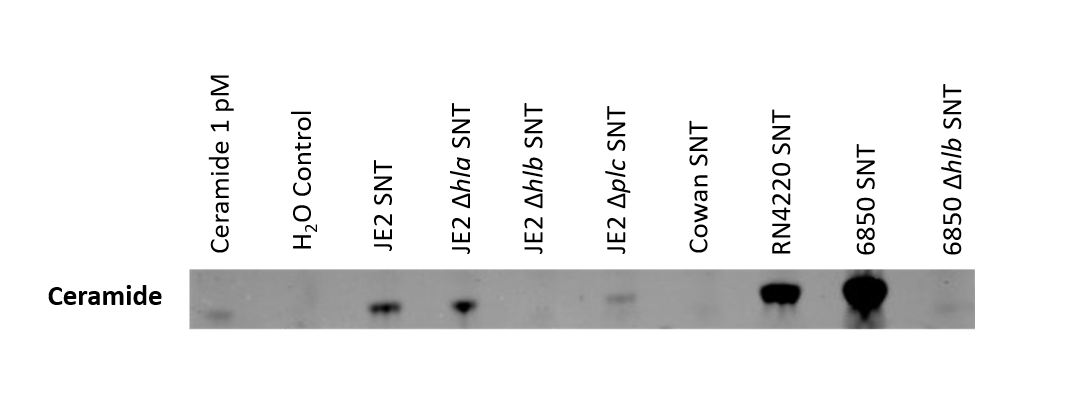

### Supplemental Figure 6

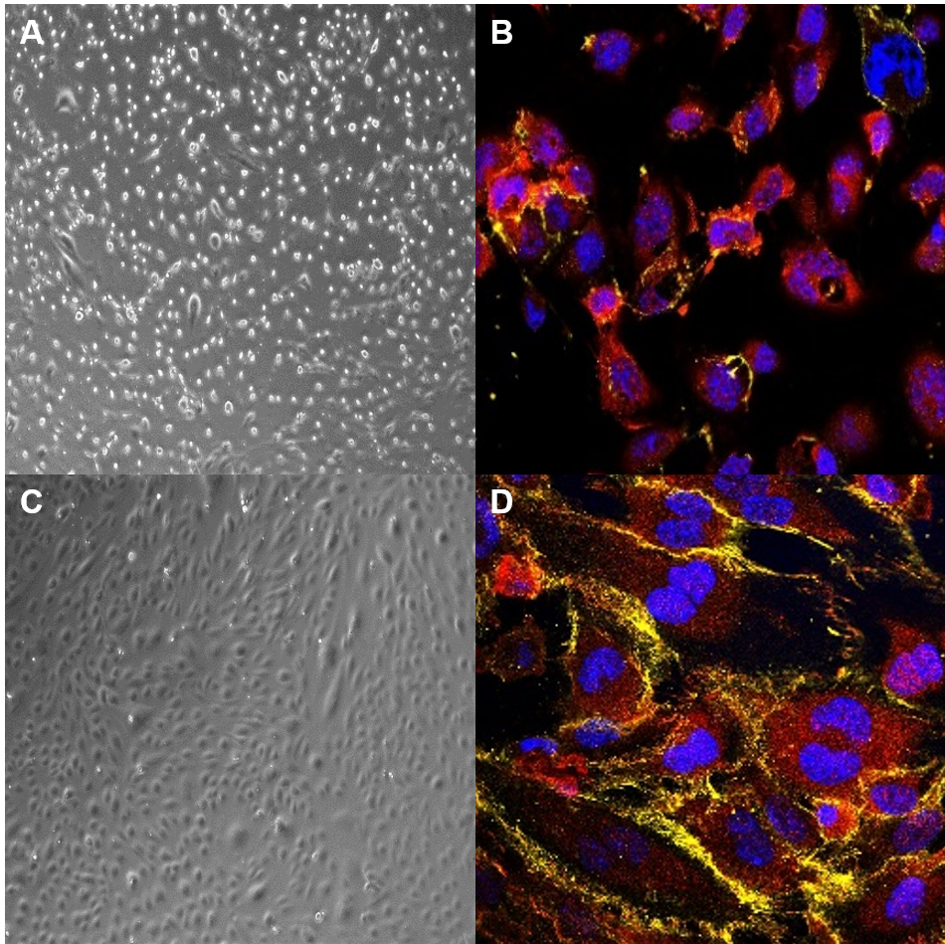

### Supplemental Figure 7

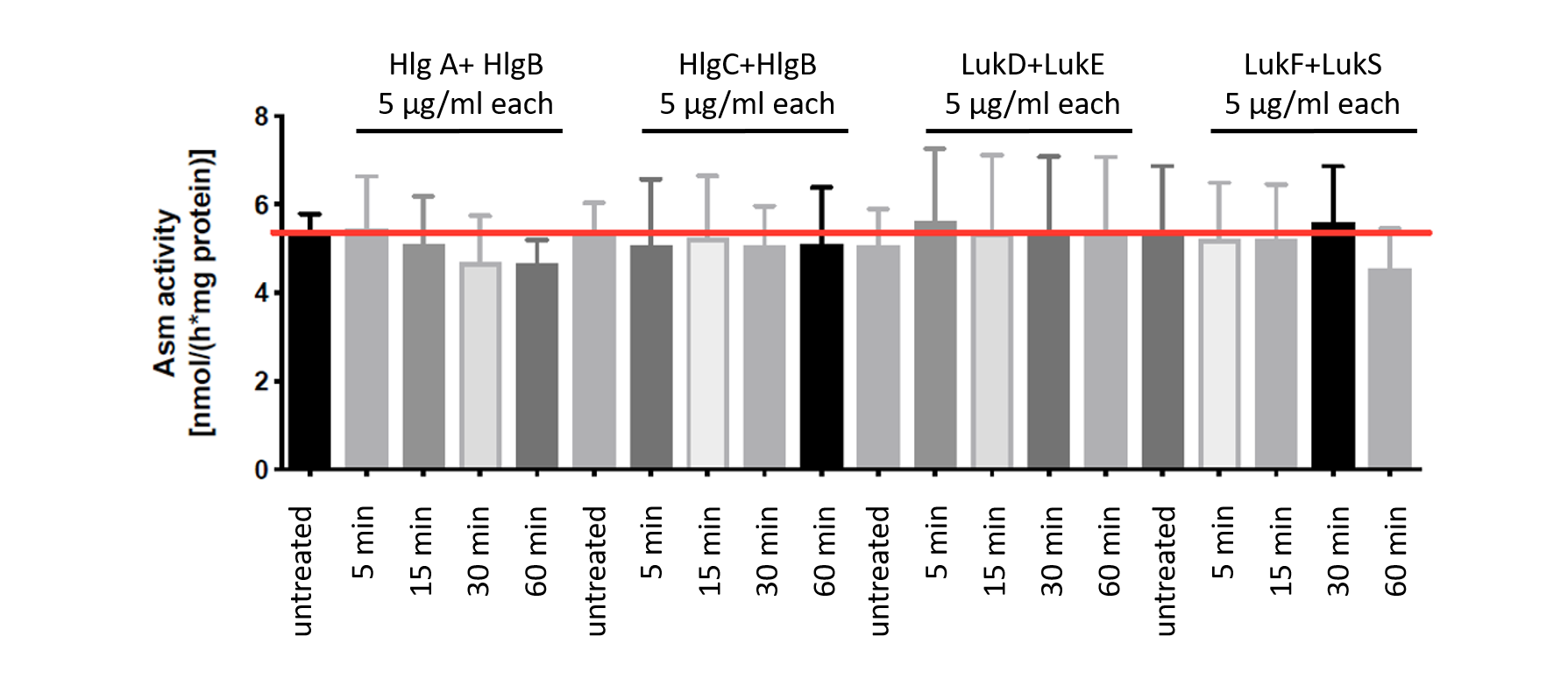
